# Seasonal weight changes in laboratory ferrets

**DOI:** 10.1101/2020.04.22.055194

**Authors:** Eleanor J. Jones, Katarina Poole, Joseph Sollini, Stephen M. Town, Jennifer K. Bizley

## Abstract

Ferrets (*Mustela putorius furo*) are a valuable animal model used in biomedical research. Ferrets undergo significant variation in body weight seasonally, affected by photoperiod, and these variations make it difficult to use weight as an indicator of health status. To overcome this requires a better understanding of these seasonal weight changes. We provide a normative weight data set for the female ferret accounting for seasonal changes, and also investigate the effect of fluid regulation on weight change. Female ferrets (n=39) underwent behavioural testing from May 2017 to August 2019 and were weighed daily while housed in an animal care facility with controlled light exposure. In the winter (October to March), animals experienced 10 hours of light and 14 hours of dark, while in summer (March to October), this contingency was reversed. Individual animals varied in their body weight from approximately 700 to 1200 g. However, weights fluctuated with light cycle, with animals losing weight in summer, and gaining weight in winter such that they fluctuated between approximately 80% and 120% of their long term average weight. Ferrets were weighed as part of their health assessment while experiencing water regulation for behavioural training. Water regulation superimposed additional weight changes on these seasonal fluctuations, with weight loss during the 5 day water regulation period being greater in summer than winter. These data establish a normative benchmark for seasonal weight variation in female ferrets that can be incorporated into the health assessment of an animal’s condition.

## Introduction

Domesticated ferrets (*Mustela putorius furo*) are valuable animal models for a wide range of biomedical research areas, including: neuroscience [1–6], drug development [7] and respiratory diseases such as Influenza and Severe Acute Respiratory Syndrome (SARS) [8,9] including the new coronavirus strain, SARS-CoV-2 [10]. In laboratory animals exposed to scientific procedures, a standard approach to monitoring health status is to measure body weight. Weight loss is a key indicator of health problems, and therefore understanding the factors that contribute to natural variation in body weight is critical for correctly monitoring an animal’s condition. Ferrets undergo significant variation in their body weight seasonally; however, there is currently no normative data available to provide a benchmark for the expected seasonal weight changes. Seasonal variations may mask or exaggerate changes in body weight due to an experimental procedure or change in health status and thus must be integrated into assessments of a ferret’s health status.

Seasonal weight changes have been demonstrated in multiple species independent of diurnality, including monkeys [11,12], raccoons [13], hamsters [14] and rodents [15]. There are a range of potential factors that elicit seasonal weight changes, but temperature and day length are key triggers, which are ultimately crucial for survival.

Ferrets are members of the mustelid family and have been domesticated from European polecats, a species which was native to western Euroasia. Seasonal weight changes have been observed in polecats and other closely related species such as mink. These weight changes are seen as adaptations to the differing energy intake and expenditure requirements of winter and summer [16,17]. In animal care facilities, daylight hours can be easily regulated and are often set at a 12-hour light cycle (12-hours ON, 12-hours OFF) or synchronised with the external environment; for example, varying from a minimum 8-hour cycle in winter (8-hours ON, 16-hours OFF) to maximum 16-hour cycle in summer (16-hours ON, 8-hours OFF) [18–21]. Variation in the photoperiod can change factors such as eating habits, coat thickness, sleep and activity levels - all of which may contribute to normal and possible abnormal weight changes. Previous research has demonstrated that ferret weights increase as hours of daylight decrease, leading to sinusoidal weight fluctuations with annual light cycle [22,23]. Contrastingly, in another study where the sleep habits of two male ferrets were tracked, light/dark schedule was shown to have no effect on their weight [24].

In addition to body weight, changes in photoperiod have also been linked to the timing of the oestrus cycle, which occurs once per year in female ferrets [20,22,25,26]. One of the first studies showed that sexual activity in ferrets increased when light duration or intensity increased [27]. Since then, further research has described ferret oestrus as seasonal and photoperiod activated [28]. The relationship between photoperiod, oestrus and body weight is unknown, but Donovan (1986) concluded that while there was not a critical weight to trigger oestrus, oestrus does require a minimum weight of around 420g.

The aim of this study is to provide data on the normative weights of female ferrets, accounting for seasonal changes over multiple years. In addition, we document changes in weight that occur due to water regulation. We hypothesized that controlled light exposure in animal care facilities would induce naturalistic fluctuations in the ferrets’ body weight.

## Methods

### Ethics Statement

All the animals in this study were maintained for the purpose of investigating the neural basis of hearing, undergoing experimental procedures that were approved by local ethical review committees (Animal Welfare and Ethical Review Board) at University College London and The Royal Veterinary College, University of London and performed under license from the UK Home Office (Project License 70/8987) and in accordance with the Animals (Scientific Procedures) Act 1986.

### Animals

The data from 39 healthy female pigmented ferrets (0.5 – 4 years) were used for this study. All animals underwent behavioural testing in psychoacoustic tasks that required regulated access to water. Water was available during twice-daily testing sessions, with supplementary wet food and/or water provided to ensure animals received a minimum of 60 ml/kg of water. Testing took place from Monday to Friday in, roughly, a three-weeks on and one week off schedule. This ensured that ferrets did not experience water regulation more than 50% of the time. When not participating in behavioural testing, animals had free access to water. During testing periods, each animal was weighed daily using digital scales (Salter, UK) prior to their morning testing session. Data was obtained from all available animals between May 2017 and August 2019 (months of participation, mean± SD: 11 months ±4.1).

Animals were housed at 15-24°C in social groups (n = 2 to 8 ferrets) and had free access to high-protein food pellets. Animals lived in enriched cages and freely exercised during daily cage cleans, with the opportunity to interact with humans, other ferrets and a variety of enrichment facilities (e.g. tunnels and balls). Our colony was comprised exclusively of female ferrets and typically contained between 25 and 30 animals.

The light cycle was changed in accordance with UK daylight savings: during ‘winter’ (October to March) ferrets were exposed to 10 hours of light and 14 hours of dark; during ‘summer’ (March to October) this was reversed to 14 hours of light and 10 hours of dark. The animal facility in which the animals were housed was windowless, and thus animals did not have access to natural light. The transition between ‘seasons’ was staggered such that timings were changed one hour per week over 4 weeks, centred on clock change for UK daylight saving time (Figure 1A).

**Figure 1:**
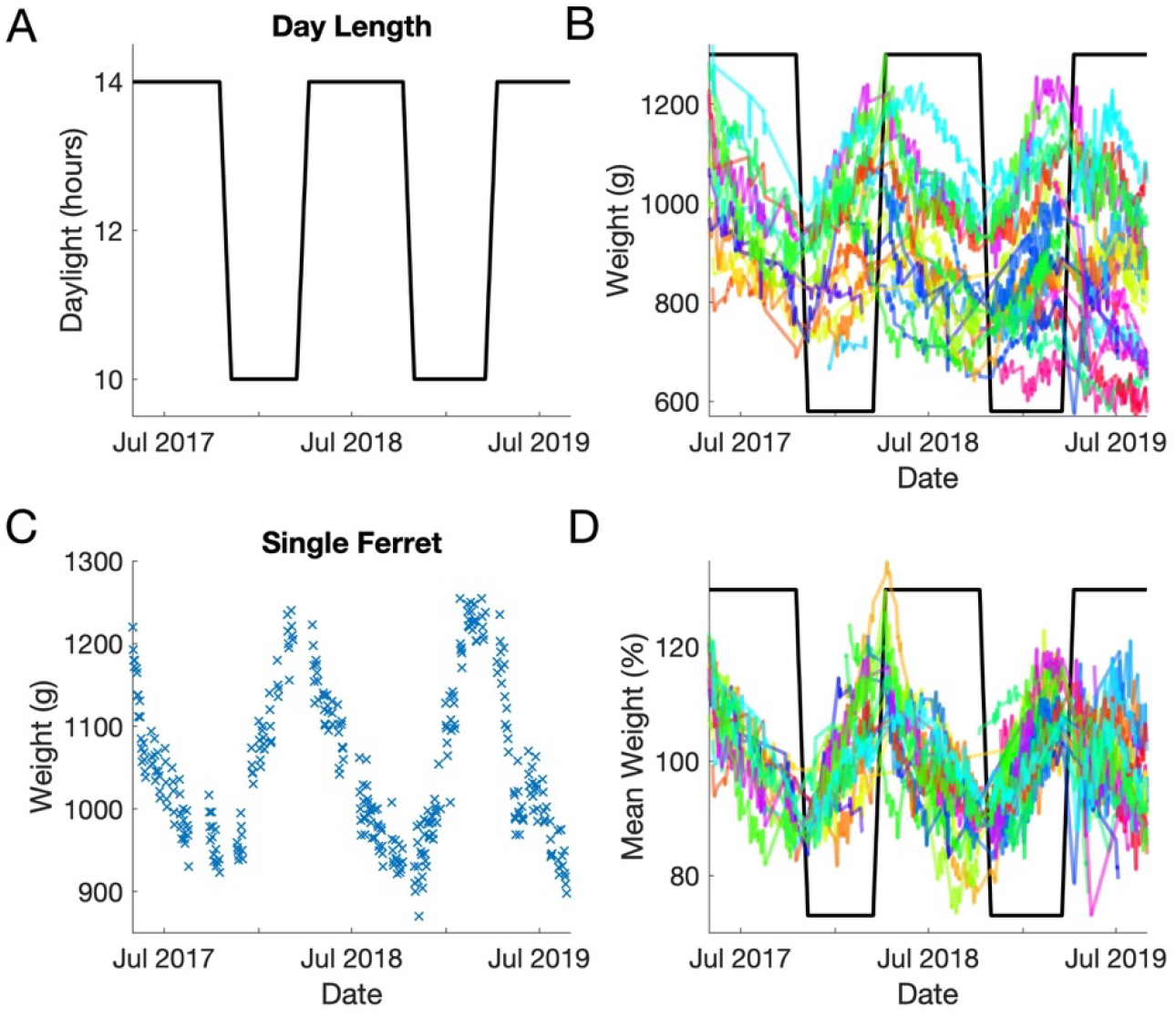
Seasonal fluctuations in body weight. **A**. Relative light hours and clock change transition periods ferrets were exposed to during the 28-month period. **B**. Weight change for a single ferret (F1606) between May 2017 and August 2019._**C**. Absolute weight of all ferrets (n=39) between May 2017 and August 2019. **D**. Seasonal variation in weight expressed as a percentage change from each ferret’s average bodyweight.

The age of each ferret was calculated from the approximate date of birth provided by the supplier (Highgate Farms, UK). Also available for each animal was the oestrus time, which was estimated from the record of each animal’s yearly hormone injection (0.5ml s.c. Proligestone, Delvosteron, Intervet). Hormone injections were given within 24-72 hours of animals exhibiting visible signs of oestrus, in order to suppress oestrus until the following spring and thus prevent life-threatening anaemia experienced by females in sustained oestrus [29].

### Data Analysis

Data was recorded and analysed in Matlab (version R2018a, MathWorks Inc, MA, USA) using custom written scripts. Weight measurements were either examined in absolute terms or relative to each animal’s long-term average (calculated from all available data). To examine day-light triggered weight changes, data from summer and winter were considered independently in terms of weeks from the transition to shorter/longer days.

ANOVAs were performed in SPSS (IBM) using Greenhouse-Geisser corrections for violations of sphericity where appropriate. Generalised linear models (GLMs) were performed using MATLAB’s ‘fitglm’ functions using a step-wise approach to fit models of increasing complexity. Model parameters were retained where an F-test indicated a significant drop in deviance upon inclusion of the term.

## Results

The weights of 39 female ferrets were recorded as part of their daily health monitoring. Weight values ranged from 553g-1350g. There was considerable variation across animals, with average weights spanning 693 to 1195g, with a population mean (±SD) of 864.8 (±119.0g). There was substantial weight variation within each animal. For example, the animal shown in Fig. 1C weighed 1240g on 5^th^ March 2018, and 870g on the 9^th^ of November that year, a change of 370g over nearly 7 months. The average standard deviation across all measurements was 64.9 (±30.3g), or equivalently, 7.45% ±3.25% of each animals’ mean weight. We next explored how variation in body weight was linked to seasonal changes and fluid regulation during behavioural testing.

### Female ferrets show significant seasonal weight variations

When weights are considered over time, cycles emerge that correlate with the seasonal light changes (Fig. 1A). The pattern of weight change for one ferret over the collection period is shown in Figure 1B. All ferrets conformed to a similar seasonal pattern of weight change with weight greatest in April (when lights were altered to their summer day length) and lowest in October (when the light cycle was switched to winter day lengths, Fig. 1C). The observed decreases in weight during the summer period and increases in weight over the winter months, resulted in sinusoidal weight fluctuations over the two-year measurement period.

To quantify the observed changes in weight with season, we divided weight measurements into ‘summer’ and ‘winter’ periods according to day length (summer = 14 hours daylight, winter = 8 hours daylight), considering time as the number of weeks since the transition to longer/shorter days. For each animal and season for which we had at least 8 weeks of data, we performed a linear regression to determine the relationship between time (in weeks) and body weight (Fig. 2A-B). In summer, there was a statistically significant relationship between time and body weight during the summer period for all animals (51 animal x transition combinations, 33 unique animals measured across one or more seasonal transition; R^2^ (mean; min to max) = 0.59; 0.07 to 0.96, p < 0.05 (49/51 p<0.001). In winter, there was a significant relationship between week and body weight for 33/37 animal-transitions (28 unique animals; R^2^ (mean; min to max), 0.74; 0.10 to 0.97 p < 0.05, 31/33 p<0.001). We, therefore, used the resulting regression coefficients (β) to determine the predicted weight change per week. We expressed weights in grams (Fig. 2C-D) and also relative to their starting weight (Fig. 2E-F). Measuring weight changes in this way allowed us to see a highly stereotyped pattern of weight loss/gain. Weight changes were negative in summer (−6.0 g/week ±5.1g/week; −0.65 % ±0.55%) and positive in winter (+8.3 g/week ± 5.2 g/week; +0.89% ±0.53%), consistent with a pattern of weight changes observed across the year.

**Figure 2:**
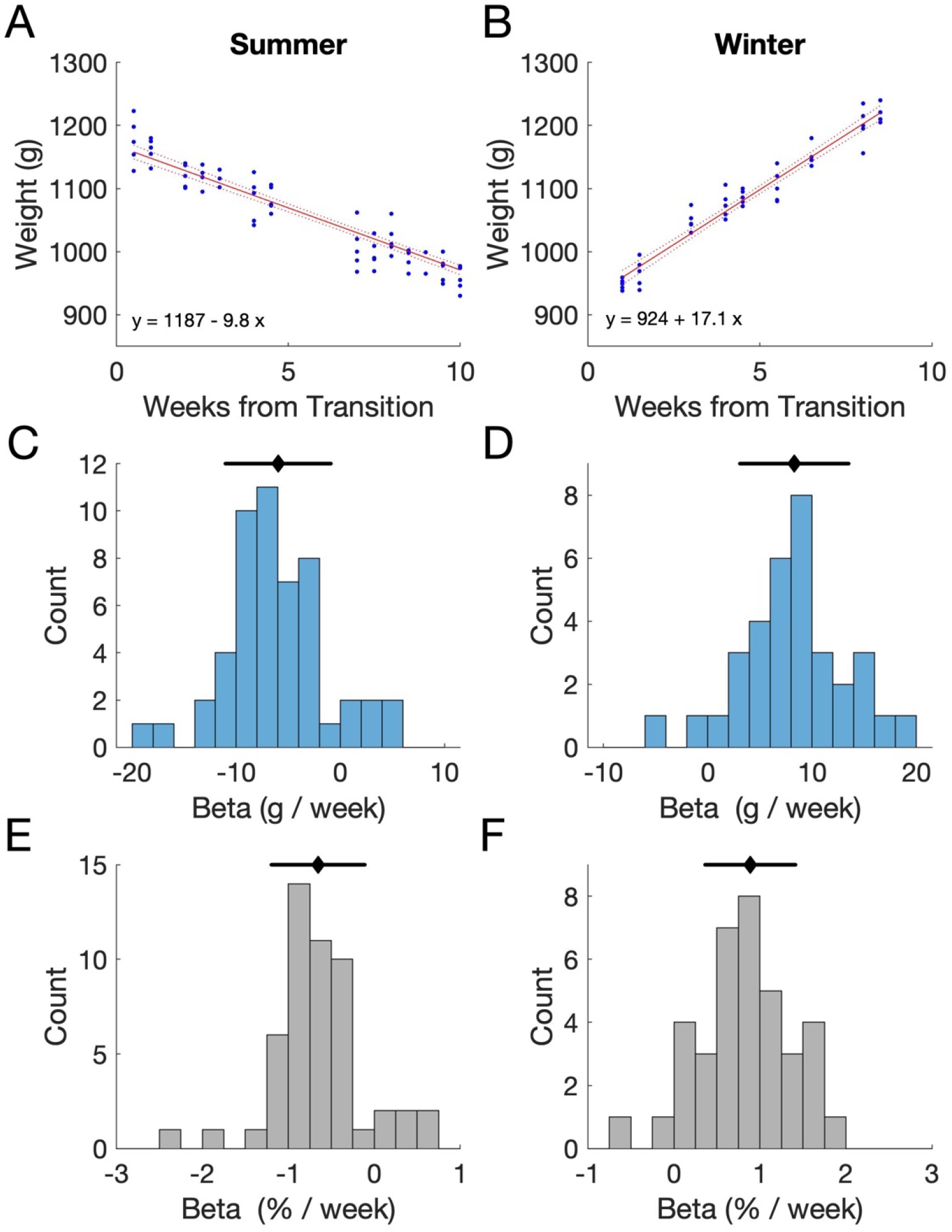
Seasonal weight changes. **A-B** Data from one animal (F1606) during summer 2018 (**A**) and winter 2017/18 (**B**). Symbols indicate individual weight measurements; plotted according to the number of weeks since the transition to summer light cycles. Line indicates the regression fit (and confidence bounds). Regression coefficients, Summer: β=-9.84 g/week or −0.94% per week (t=-26.8, p<0.001); Winter: β=17.01 g/week or 1.66% per week (t=27.8, p<0.001). **C-D** Regression coefficients for all unique animal-transition combinations between 2017-2019 during summer (**C**, n=51 ferret x transition combinations, 33 unique animals) and winter (**D**, n=37 ferret x transition combinations, 28 unique animals). Black lines indicate the mean and standard deviation. **E-F** Regression coefficients from C-D expressed as percentage of long-term mean body weight.

### Water regulation is associated with predictable weight variation

We next consider the impact of fluid regulation on body weight. The ferret is a popular model for neuroscience research as animals can be readily trained in a variety of complex behavioural tasks using water as a positive reward [30–32]. The animals that formed this dataset performed psychoacoustic tasks and were weighed as part of their daily health monitoring while on water regulation. Water regulation typically took place over a 5-day cycle, with water being removed from the home cage from Sunday night until Friday afternoon. We sought to quantify the impact of regulation on body weight, and whether there was any interaction with season changes reported above.

We divided data into summer and winter periods and compared day of water regulation to bodyweight in absolute terms (Fig. 3A) or relative to each animal’s long-term mean summer or winter weight (Fig. 3B). Both metrics show that weight declined with day on water regulation, although the trends differed between summer and winter. In the winter weight loss reached a stable baseline, whereas in the summer weight loss appeared more linear. These results were confirmed statistically using a two-way repeated measures ANOVA to analyse absolute body weight, where we found a main effect of day (F_4, 116_ = 59.4, p<0.001) and interaction between day and season (F_4, 116_ = 3.51, p = 0.023). Similar results were also found when analysing relative body weight, where again both main effect of day (F_4, 116_ = 59.1, p < 0.001) and interaction between day and season (F_4, 116_ =3.93, p = 0.014) were significant. In neither analysis was the effect of season significant by itself.

**Figure 3:**
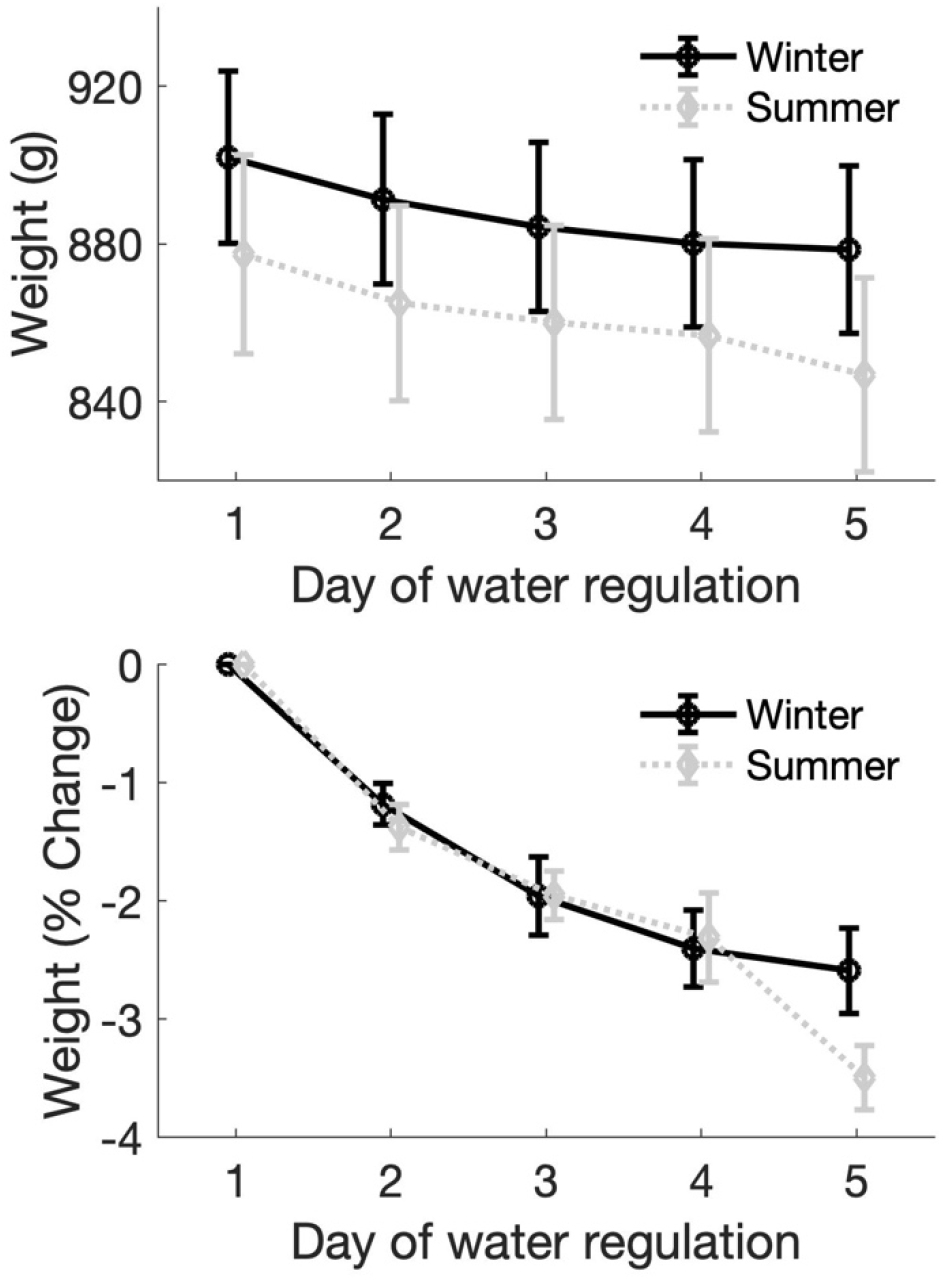
Weight changes resulting from water regulation: **A-B** Water regulation related weight changes for 30 animals in which both summer and winter data were available. **A**; mean± SEM weight in g for each day of water regulation for summer and winter periods. **B** mean ± SEM weight change expressed as a percentage change from the weight measurement made on a Monday morning (day 1).

### Modelling the contribution of season and fluid regulation to body weight

To determine the relative contribution of season and water regulation duration on body weight, we fitted General Linear Models to weight data, using the following predictors: starting weight (the first measurement of weight made after the light transition, this data point was excluded from the modelling), the number of weeks since light transition, and the day of water regulation. We again considered summer and winter data separately, and for each, found the best fitting model using a significant drop in deviance as the criterion for including parameters (using the F statistic to compare models, p<0.05). In each case the best fitting mode retained each of the three main effects (week since transition, day of water regulation and starting weight) as well as the two-way interaction between the week and starting weight (see table 1). To illustrate the key features of these models, we used the fitted models to simulate the changes in weight that would occur over a 20-week period in summer and winter for animals of 750g and 900g (Fig. 4).

**Figure 4:**
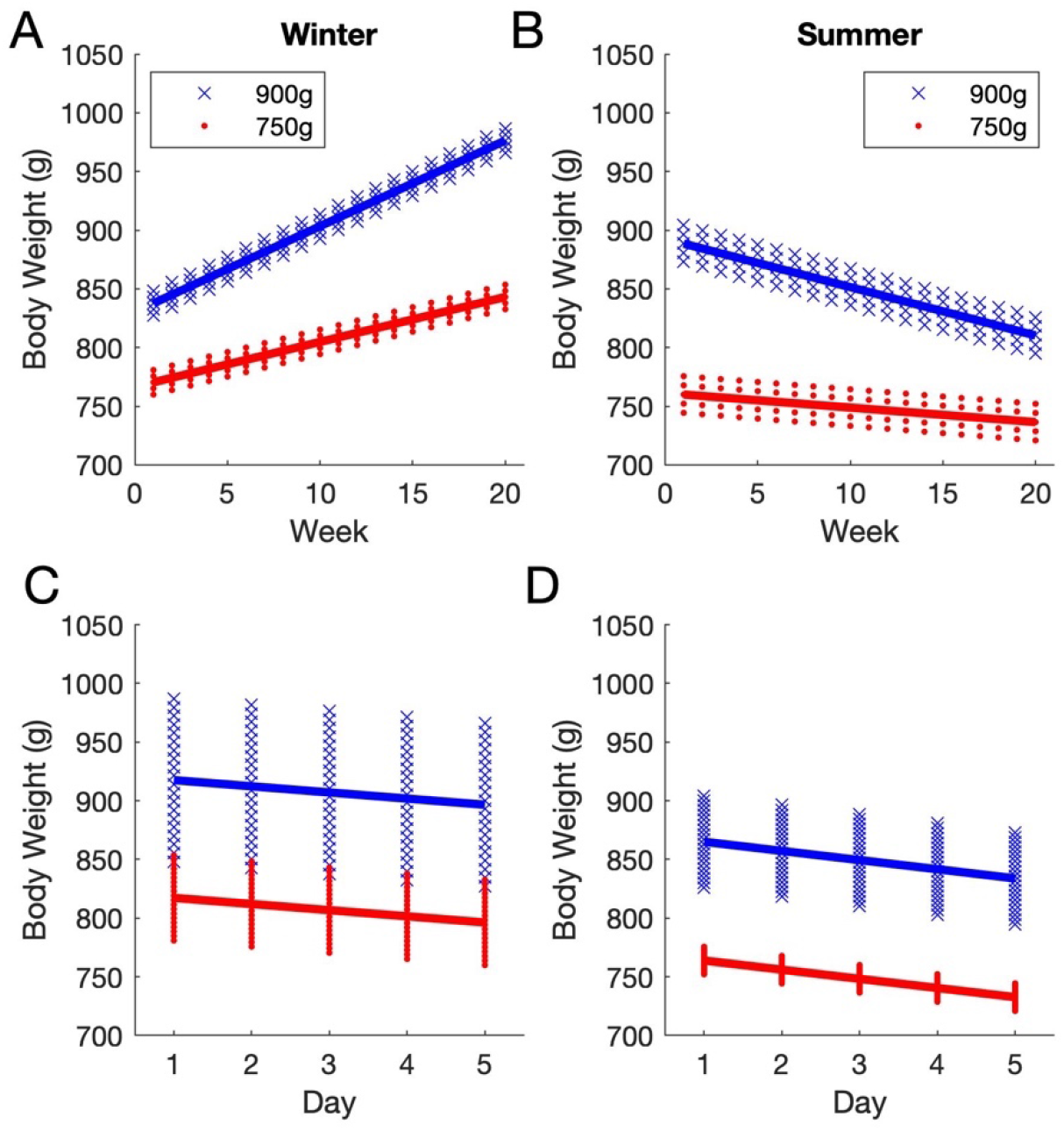
Predicted weight changes. **A**-**B**, predicted weight changes over a 20-week period for two simulated ferrets of 750g (red) and 900g (blue) in winter (**A**) and summer (**B**). Predictions were generated for each simulated animal for 20 weeks at each light duration, for 5 days of water regulation within each week (i.e. 100 values per season). **C-D** Predicted within-week changes during winter (**C**) and summer (**D**).

**Table 1:**
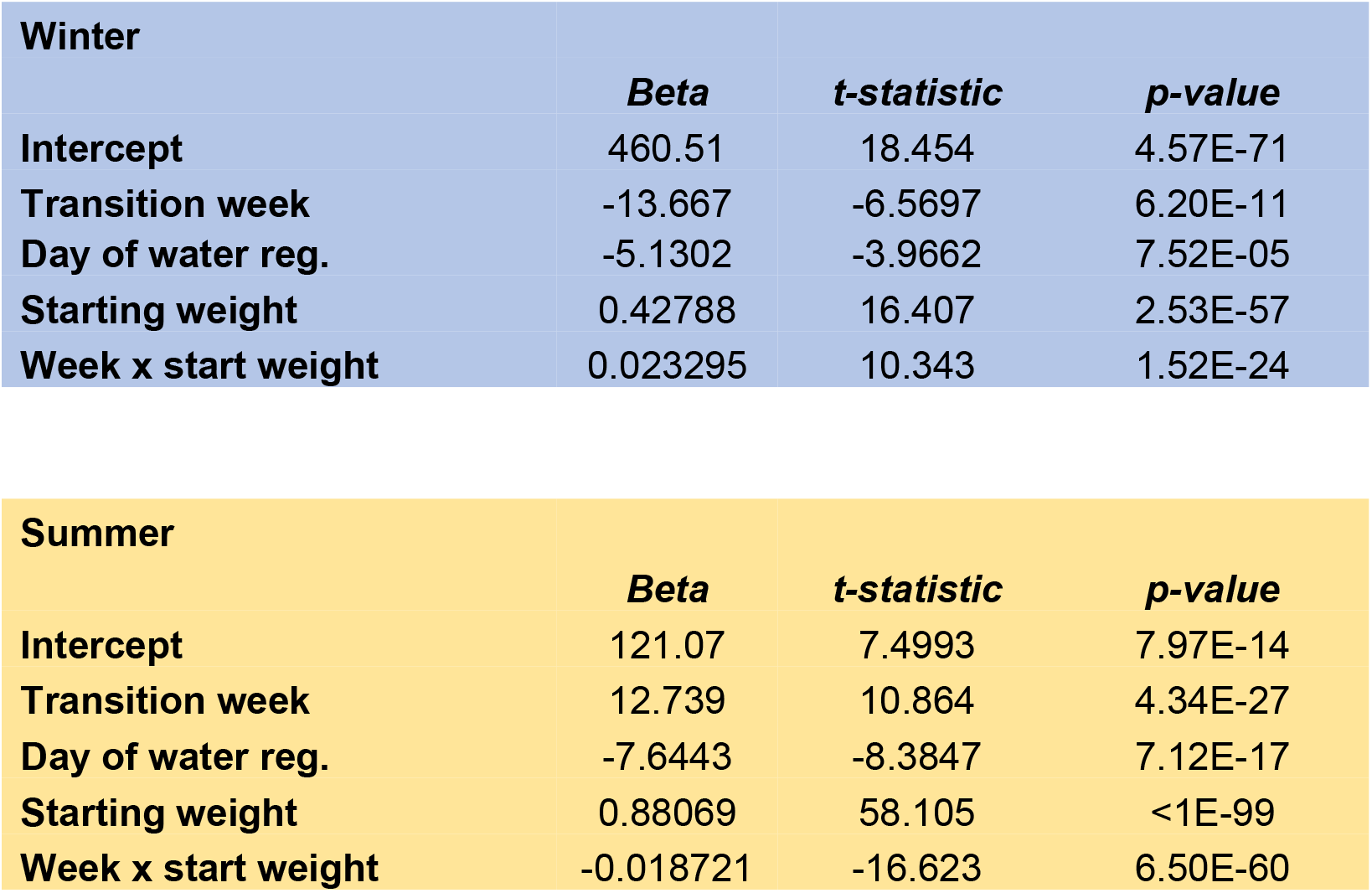
GLM results for summer and winter data. Only significant predictors for which there was a significant reduction in model deviance are shown. For both summer and winter weights the resulting model retained the starting weight of the animal, the week of transition, and the day of water regulation, as well as a significant start weight * day of regulation interaction.

| Winter |  |  |  |
| --- | --- | --- | --- |
|  | Beta | t-statistic | p-value |
| Intercept | 460.51 | 18.454 | 4.57E-71 |
| Transition week | -13.667 | -6.5697 | 6.20E-11 |
| **Day of water reg.** | -5.1302 | -3.9662 | 7.52E-05 |
| **Starting weight** | 0.42788 | 16.407 | 2.53E-57 |
| **Week x start weight** | 0.023295 | 10.343 | 1.52E-24 |

**Table 1:** GLM results for summer and winter data.
| **Summer** |  |  |  |
| --- | --- | --- | --- |
|  | **Beta** | **t-statistic** | **p-value** |
| **Intercept** | 121.07 | 7.4993 | 7.97E-14 |
| **Transition week** | 12.739 | 10.864 | 4.34E-27 |
| **Day of water reg.** | -7.6443 | -8.3847 | 7.12E-17 |
| **Starting weight** | 0.88069 | 58.105 | <1E-99 |
| **Week x start weight** | -0.018721 | -16.623 | 6.50E-60 |

While our models could recapitulate key trends in the data, and for some animals the predicted and observed weights were closely aligned (Figure 5A), others were much less well predicted (e.g. Figure 5B). Moreover, to be useful as a diagnostic measure of healthy or abnormal weight changes, the model should be able to estimate an expected weight of an animal given factors such as the season and its starting weight. To test this, and determine whether this model had any utility as a diagnostic measure, we applied the regression model obtained by fitting data to all animals (above) to data from all but one animal, excluding each animal in turn. We used the resulting model to predict weight measurements (and their 99% confidence intervals) for the left-out animal’s data and then compare predicted and actual weight measurements.

**Figure 5:**
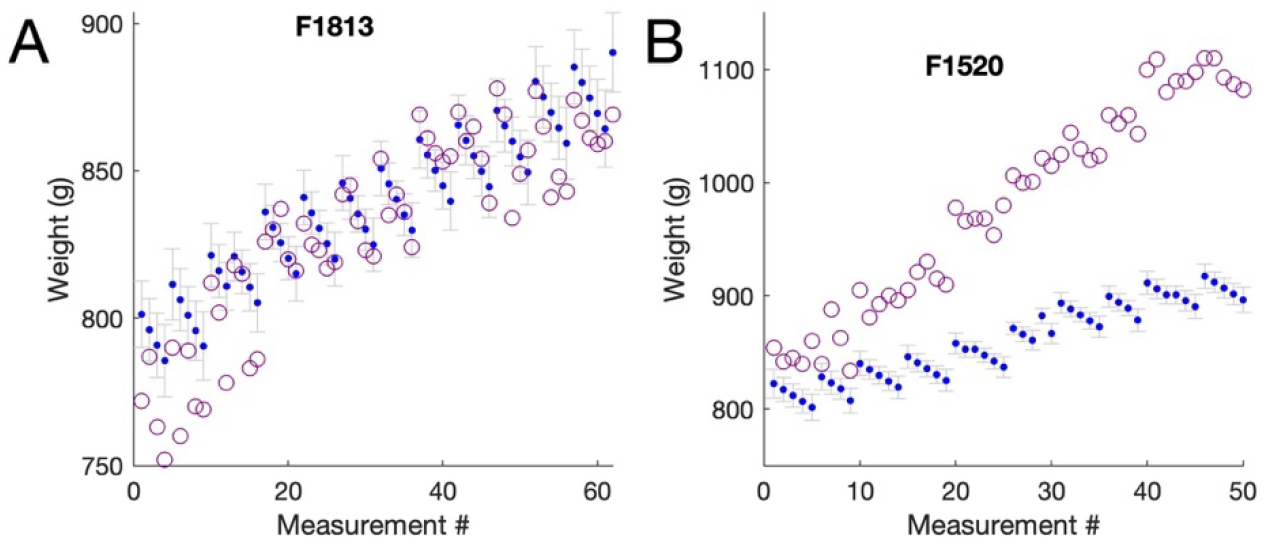
Comparisons of observed and predicted values for two ferrets (A, F1830, winter, B, F1520, winter) selected as two examples of animals whose observed weight values were relatively well fit (A) and poorly fit (B) by data predicted from a GLM modelling expected weight values based on starting weight, number of weeks since light transition and day of water regulation. Open data points show the observed weight values, dots and errorbars show the 99% confidence intervals of the model predictions.

We found that that although predicted and observed weights were correlated (significant correlations were found for 29/30 animals in summer with r values (mean±SD) r=0.69±0.19, 24/32 animals in winter; r=0.72±0.33), generally the ability to predict ‘held-out’ animals was poor. Very few of our observed weight measurements fell within the predicted confidence limits (~10% of all data). Examination of the observed and predicted values revealed that while the models predicted the key trends, they poorly captured the extent of the variability occurring within individual animals over time. In Figure 6, four example animals are shown – the first two animals show what is typical of most of our animals which is that the model captures the trend but not the details, whereas the second two animals show very different patterns of weight loss which, while quite different from most animals is highly consistent within that individual.

**Figure 6:**
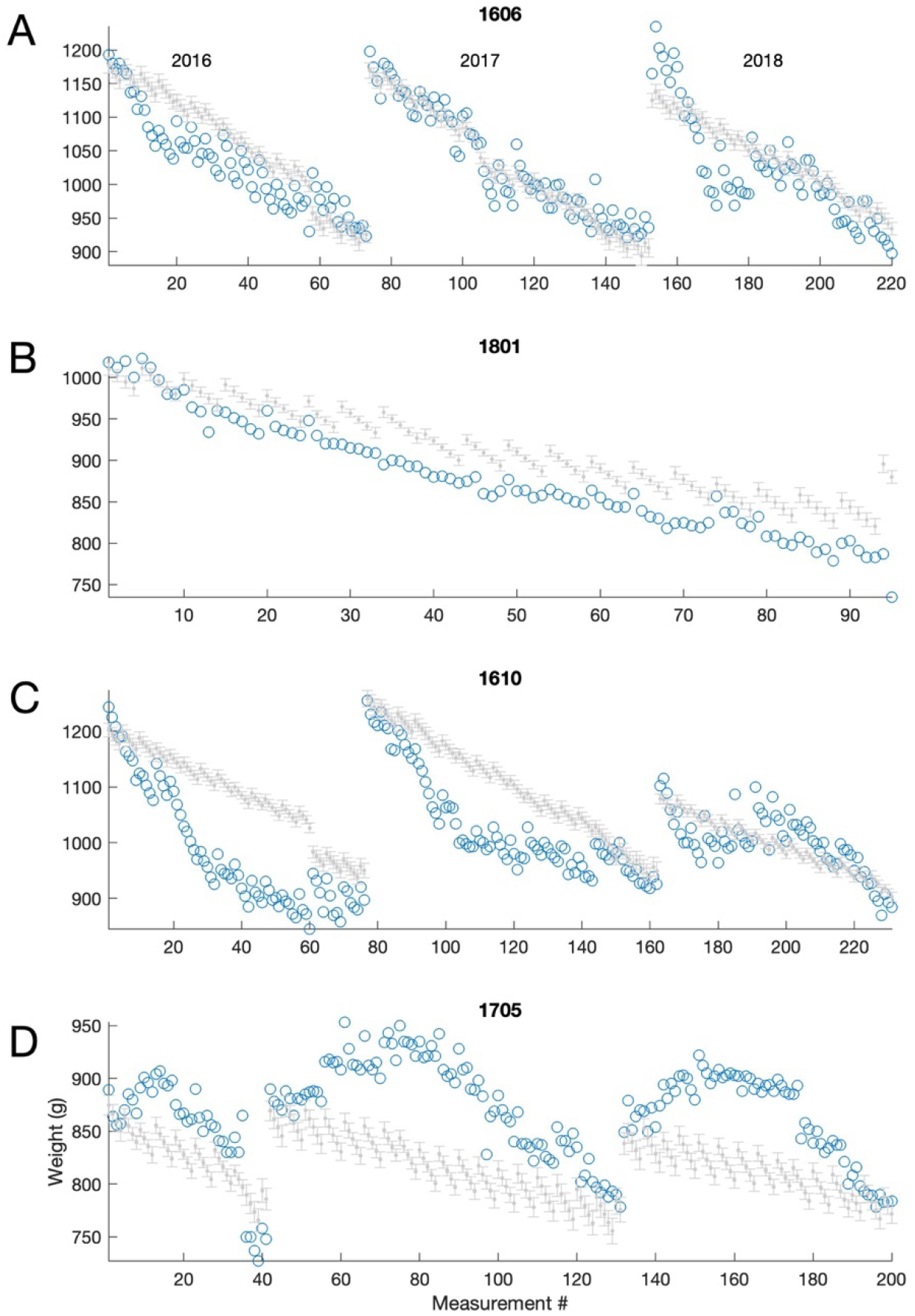
Comparisons of observed (open symbols) and predicted values (gray error bars indicating the 99% confidence interval of the fit) for four animals. For each animal all available data is shown, which includes data from each of three summers 2016-2018, for animals in A,C,D, and a single year (2018) in animal F1801, shown in B. The animals in A and B show the typical near-linear pattern of weight loss over time, the shape of which is well captured by the predictions. However, rarely are the observed values within the predicted range by the model. In animal A, the fluctuation in weight due to water regulation is underestimated, and in one year weight loss was more rapid than the model predicted, while in B, the animal varies very little while on water regulation and therefore the fluctuations are over estimated. For this animal the model also underestimates the gradient of weight loss through the summer. Most animals looked like those in A and B, however a few animals had substantially different patterns which were highly consistent within each animal from year to year. Both animals in C and D show a pattern of weight loss which is not linear (and therefore unsurprisingly poorly modelled). Animal F1610 (C) shows a rapid weight loss that stabilises at a baseline, whereas Animal F1705 continues to gain weight after the weight change, before beginning to lose weight for the remainder of the season.

Given the observation that weight loss/gain with season could be fit as a linear model (Figure 2E,F), but that there was significant inter-animal variation we adapted our modelling approach to address three questions about the changes in body weight between weeks that occurred with water regulation that were common across animals: First, was the change in weight between weeks constant, or did it vary through the season? Second, does the change in weight between weeks, expressed as a proportion of the animal’s weight, depend on the size of the animal? Third, does some of the additional variability we observed relate to whether animals have been on water regulation the previous week? To address this, we first considered the change in weight that occurred from each week (week *i*) to the next (week *i+1*) (using only the body weight measurements from the first day on water regulation, i.e. Monday mornings), expressing this as a % change in weight relative to bodyweight measured in week *i*. We then fitted a GLM to these % values, using three predictors: (1) the number of weeks since seasonal transition (i.e. summer to winter), (2) the animal’s mean weight, and (3) whether the animal had been on water regulation the previous week (week *i-1*). As before, data for summer and winter were modelled separately.

When modelling weight changes in the winter the intercept was significant (beta = 1.38, table 2), suggesting that typically animals gained 1.38% of their body weight weekly in winter, and the week was also a significant predictor (beta = −0.065) indicating that animals initially gained weight more rapidly. In summer the intercept was again significant (beta = −0.73) and the only other significant predictor was whether the animal had been on regulation the previous week (beta = −0.82, indicating that animals who had access to free water in the previous week lost more weight than those who had not had access to free water). In summary therefore, it is possible to estimate typical patterns of weight loss / gain that should occur over the course of a week but data such as that shown in Figure 6C,D demonstrate that each animal must really be assessed individually (ideally in comparison to its own historic data) in the context of other factors.

**Table 2.** Final models for estimating the % change in body weight from week *n* to *n+1*. The number of weeks since transition, the animals mean body weight and whether the animal had been water regulated in week *n-1* were considered as factors, with only the number of weeks since transition being predictive for the summer data, and whether the animal had been on water regulation or not the previous week influencing the summer data.

| Winter |  |  |  |
| --- | --- | --- | --- |
|  | *Beta* | *t-statistic* | *p-value* |
| Intercept | 1.3778 | 6.3158 | 6.63E-10 |
| Free water previous week | -0.065161 | -3.3247 | 9.60E-04 |

Table 2
| Summer |  |  |  |
| --- | --- | --- | --- |
|  | *Beta* | *t-statistic* | *p-value* |
| Intercept | -0.73 | -9.8469 | 1.65E-21 |
| Free water previous week | -0.82439 | 4.1028 | 4.56E-05 |

In addition to body weight data, we also had the timing of oestrus for each animal (69 measurements, 39 unique animals). Oestrus varied from 2 to 8 weeks after the first change in light cycle length in the spring. Timing did not vary significantly across the three years (Kruskalwallis test, p=0.54) with the average value being 5.7±1.4 weeks after the first light change in the spring (Figure 7). Unfortunately, the number of missing weight values obtained on the week of light change and the week of oestrus precluded meaningful statistical analysis of the relationship between weight change and oestrus.

**Figure 7.**
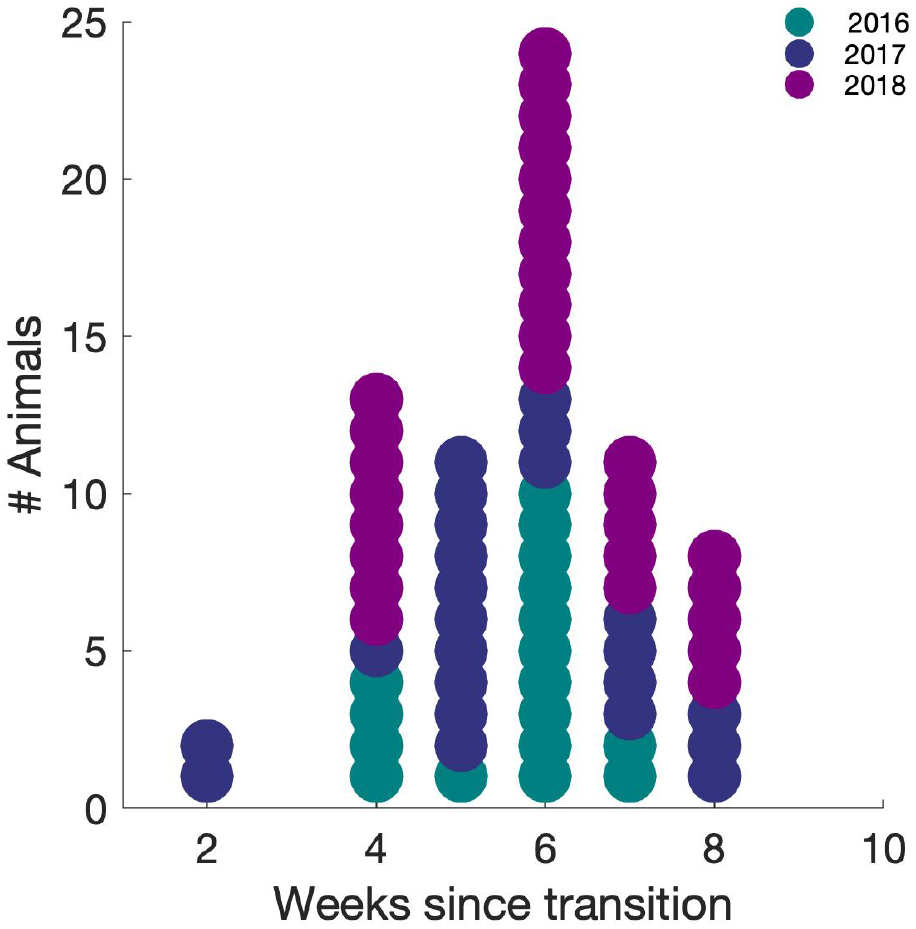
Timing of oestrus relative to daylight change. Dot histogram with each dot representing the timing of oestrus in a single animal in 2016. 2017 and 2018.

## Discussion

Here we provide a normative weight dataset for the healthy female ferret and demonstrate that ferrets show predictable and stereotypical seasonal fluctuations in weight, with most animals gaining around 0.89% of their average body weight per week in winter and losing around 0.65% of their weight per week in summer. Superimposed upon these seasonal fluctuations, water regulation also causes highly stereotyped changes in body weight.

The observed seasonal changes in weight imposed large fluctuations in body weight with animals typically being roughly 15% heavier in winter, and 15% lighter in summer (i.e. a variation of as much as 30% of their mean weight). This pattern of seasonal weight change demonstrated by our ferrets follows previously observed changes [22]. The range of mean weights that we observed across our population were in keeping with previously reported data [20,29,33]. There are many possible physiological contributing factors to this seasonal weight loss including coat changes [22,26], fat storage, hormone levels such as melatonin [26] and activity levels which in wild animals are critical for survival [34].

Water regulation imposed an additional pattern of weight changes on animals; weight was lost over the week in both summer and winter. On average, between Monday morning (when water bottles had been removed the previous evening) and Friday morning animals lost around 3% of their body weight in winter, and 4% in summer. Depending on diet, water consumption for a ferret can be up to 100ml/day [29] and we ensured ferrets received 60ml/kg of water each day of water restriction (which is the amount that animals maintained on laboratory ferret diet, with free access to water, typically consume in a 24 hour period). Since the key contributor to weight loss in water regulated animals is thought to be reluctance to eat dry food (rather than dehydration per se) providing animals with water combined with ground pellet diet to form a mash [8] appears to be successful at ensuring weight loss does not exceed more than a few percent. Food and water restriction are common methods used as motivation to train many laboratory animals including ferrets, rats and mice in tasks for research [35] and weight loss is a key marker of health status. Understanding how seasonal fluctuations interact with these effects is therefore important to refine health assessment and ensure the highest standards of animal welfare.

In addition to body weight data, we also had the timing of oestrus for each animal (69 measurements, 39 unique animals). Oestrus varied from 2 to 8 weeks after the first change in light cycle length in the spring with an average value of 5.7±1.4 weeks. Previous research has observed that the cycle of changes in light duration were responsible for initiating oestrus and weight changes but whether there is a causal relationship between the two remains unknown [20]. Further research is required to directly assess whether it is the day light change itself, weight changes induced by day light change, or an interaction between the two factors that induce oestrus.

The animals used in this study were all classed as young to middle aged [29] with the oldest animal being 4 years old. Ferret weights also vary across the lifespan, which is on average between 6-8 years [29]. At birth, female ferrets weigh 6-12g and grow rapidly to 550-700g at 10-12 weeks and 600-950g at approximately 16 weeks (i.e. adulthood) [8,20,23,29]. Ferrets are defined as old after the age of 3-4 years and are at greater risk of geriatric diseases but also natural weight loss [29,36].

We do not have sufficient repeated data from older animals to determine how ageing interacts with seasonal weight changes. Greater weight loss is observed in aged ferrets (>4 years old) during, and in recovery from, illness [37]. With weight loss an indicator of possible disease, accounting for the age of the animal is important for contextual assessment of health status, and so further research is required to quantify weight changes during ageing and precautionary close observation of weight loss in older animals would be justified.

While the seasonal and water-regulation induced changes in weight were highly stereotyped within an animal, and had common features across animals, we were unable to generate a simple statistical model that could accurately predict the expected weight changes. Pulling together our findings we would expect that animals should gain weight in winter, with an initial increase of around roughly 1.4%/week, declining to 0.7%/week after 10 weeks and 0.1%/week after 20 weeks. In contrast, expected weight loss in the summer was roughly linear, with animals losing roughly 0.7%/week, except in weeks which were preceded by access to free water in which weight loss was around 1.5%. Superimposed upon these weekly changes are daily fluctuations in body weight that result from fluid regulation which are typically of the order of 3-4% from a Monday morning to Friday morming. These data therefore establish some normative benchmarks for seasonal weight variation in female ferrets that can be incorporated along with other indicators of well-being into the assessment of an animal’s overall condition.

## Acknowledgements

This research was funded by a Royal Society & Wellcome Trust fellowship to JKB (098418/Z/12/Z). We are grateful for the technical assistance of the Royal Veterinary College in maintaining our ferret colony.

